# Transmission Bottleneck Size Estimation from Pathogen Deep-Sequencing Data, with an Application to Human Influenza A Virus

**DOI:** 10.1101/101790

**Authors:** Ashley Sobel Leonard, Daniel Weissman, Benjamin Greenbaum, Elodie Ghedin, Katia Koelle

**Affiliations:** Department of Biology, Duke University, Durham, NC 27701; Department of Physics, Emory University, Atlanta, GA 30322; Tisch Cancer Institute, Departments of Medicine, Oncological Sciences, and Pathology, Icahn School of Medicine at Mount Sinai, New York, NY, 10029.; Center for Genomics & Systems Biology, Department of Biology, and College of Global Public Health, New York University, 12 Waverly Place, New York, NY 10003

## Abstract

The bottleneck governing infectious disease transmission describes the size of the pathogen population transferred from a donor to a recipient host. Accurate quantification of the bottleneck size is of particular importance for rapidly evolving pathogens such as influenza virus, as narrow bottlenecks would limit the extent of transferred viral genetic diversity and, thus, have the potential to slow the rate of viral adaptation. Previous studies have estimated the transmission bottleneck size governing viral transmission through statistical analyses of variants identified in pathogen sequencing data. The methods used by these studies, however, did not account for variant calling thresholds and stochastic dynamics of the viral population within recipient hosts. Because these factors can skew bottleneck size estimates, we here introduce a new method for inferring transmission bottleneck sizes that explicitly takes these factors into account. We compare our method, based on beta-binomial sampling, with existing methods in the literature for their ability to recover the transmission bottleneck size of a simulated dataset. This comparison demonstrates that the beta-binomial sampling method is best able to accurately infer the simulated bottleneck size. We then apply our method to a recently published dataset of influenza A H1N1p and H3N2 infections, for which viral deep sequencing data from inferred donor-recipient transmission pairs are available. Our results indicate that transmission bottleneck sizes across transmission pairs are variable, yet that there is no significant difference in the overall bottleneck sizes inferred for H1N1p and H3N2. The mean bottleneck size for influenza virus in this study, considering all transmission pairs, was *N*_b_ = 196 (95% confidence interval 66-392) virions. While this estimate is consistent with previous bottleneck size estimates for this dataset, it is considerably higher than the bottleneck sizes estimated for influenza from other datasets.

**Author Summary:** The transmission bottleneck size describes the size of the pathogen population transferred from the donor to recipient host at the onset of infection and is a key factor in determining the rate at which a pathogen can adapt within a host population. Recent advances in sequencing technology have enabled the bottleneck size to be estimated from pathogen sequence data, though there is not yet a consensus on the statistical method to use. In this study, we introduce a new approach for inferring the transmission bottleneck size from sequencing data that accounts for the criteria used to identify sequence variants and stochasticity in pathogen replication dynamics. We show that the failure to account for these factors may lead to underestimation of the transmission bottleneck size. We apply this method to a previous dataset of human influenza A infections, showing that transmission is governed by a loose transmission bottleneck and that the bottleneck size is highly variable across transmission events. This work advances our understanding of the bottleneck size governing influenza infection and introduces a method for estimating the bottleneck size that can be applied to other rapidly evolving RNA viruses, such as norovirus and RSV.

## Introduction

Infectious disease transmission relies on the transfer of a pathogenic organism from one host to another. This transfer is characterized by a transmission bottleneck, defined as the size of the founding pathogen population in the recipient host. Accurate quantification of transmission bottleneck sizes for pathogenic organisms is critical for several reasons. First, bottleneck sizes impact levels of genetic diversity in recipient hosts, and thereby impact the rate at which pathogens can adapt to host populations, with smaller bottleneck sizes slowing rates of adaptation [1,2]. Second, when cooperative interactions occur within a pathogen population (e.g., [3,4]), or when viral complementation and cellular coinfection are critical for producing viral progeny (e.g., [5]), bottleneck sizes will necessarily impact initial pathogen replication rates, with larger bottleneck sizes enabling the occurrence of these interactions and thus facilitating within-host replication. Finally, transmission bottleneck sizes impact the ability to accurately reconstruct who-infected-whom during an ongoing epidemic [6], such that estimation of the transmission bottleneck size can point to cases which may be problematic, and for which a certain class of phylodynamic inference methods (such as [7]) might be particularly useful.

The transmission bottleneck size has been estimated for a number of pathogenic organisms, including pathogens of plants [8–13] and animals [14–22]. While these estimates have relied on the distribution of pathogen types in the infection recipients, as determined by molecular and phenotypic markers or Sanger sequencing of the pathogen population in donor and recipient hosts, deep sequencing data have recently started to be used to gauge transmission bottleneck sizes [23–29]. Some of these studies have characterized the general magnitude of transmission bottlenecks size, with results indicating that narrow, selective bottlenecks tend to govern the transmission dynamics of viral pathogens that are ill-adapted to their recipient hosts [24–26]. Studies that have instead gauged transmission bottleneck sizes of well-adapted viral pathogens using deep sequencing data have, in contrast, generally found that they tend to be loose, with many virions initiating infection [23,28,29]. While many of these studies focus on assessing how “loose” or “narrow” a transmission bottleneck is, other studies have attempted to more quantitatively estimate transmission bottleneck sizes. One approach relied on the use of barcoded influenza virus during experimental transmission studies in small mammals, with results indicating that the route of transmission greatly impacts the size of the bottleneck [27].

In natural infections, it is not feasible to rely on barcoded or otherwise marked pathogens. In these cases, statistical approaches have therefore instead been used to quantify bottleneck sizes [28,30]. Two studies have used the Kullback-Leibler divergence index (developed in [30]) to estimate the viral effective population size initiating infection from deep sequencing data [28,30]. One of these studies quantified the transmission effective population size for ebola in human-to-human infections [30]. The other quantified this transmission effective population size for human influenza A viruses [28]. A second statistical approach used by [28] makes use of a single-generation population genetic Wright-Fisher model to estimate the effective viral population size initiating infection. While this approach similarly showed that the effective population size following influenza virus transmission in natural human-to-human infection is large, this model yielded quantitatively different results from the Kullback-Leibler approach. Further, in both of these studies, it is not clear how the effective population size relates to the transmission bottleneck size. It is worth noting, however, that the effective population size is generally considered to be an underestimate of the true population size as it represents the minimum population size necessary to establish observed levels of genetic diversity.

Both of these approaches [28,30] analyze only variants that are identified as present in both the donor and the recipient. But the absence of a donor variant in a recipient host is also informative, and ignoring such missing variants can significantly bias transmission bottleneck size estimates. Another limitation of both approaches is that they do not consider the effect stochastic dynamics early in infection may have on variant frequencies in the recipient. To address these concerns, we here introduce a new method for estimating the transmission bottleneck size of pathogens. This method accounts for stochastic dynamics occurring during viral replication in the recipient and further accounts for variant calling thresholds that are used in calling a variant present or absent in a sample. We refer to this method as the beta-binomial sampling method, based upon the method’s derived likelihood expression. Using a simulated dataset, we compare the beta-binomial sampling method to two methods of bottleneck size inference that are present (in some form) in the current literature: the presence/absence method and the binomial sampling method. This comparison demonstrates that the beta-binomial sampling method is able to recover the true bottleneck size of the simulated dataset, whereas the 2 other methods infer biased estimates by failing to account for variant calling thresholds or stochastic dynamics in the recipient host. Finally, we apply the beta-binomial sampling method to an existing next generation sequencing dataset of influenza A virus infections to estimate the transmission bottleneck size in natural human-to-human flu transmission.

## Models

Figure 1 provides a schematic of the data that are used for inferring transmission bottleneck sizes in the approaches we consider in this study. Deep sequencing data consist of short reads at various sites in the genome, obtained from both the infected donor and the recipient at, generally, a single time point for each individual. The short read data are used to identify viral variants in the donor and recipient hosts. Comparison of these variants’ frequencies across donor-recipient transmission pairs allows us to infer the transmission bottleneck size (*N*_b_), the number of virions comprising the founding viral population at the onset of infection in the recipient host. We specifically define *N*_b_ as the number virions that successfully establish lineages that persist to sampling; there may be additional virions that transiently replicate in the recipient host but quickly die out.

**Figure 1.**
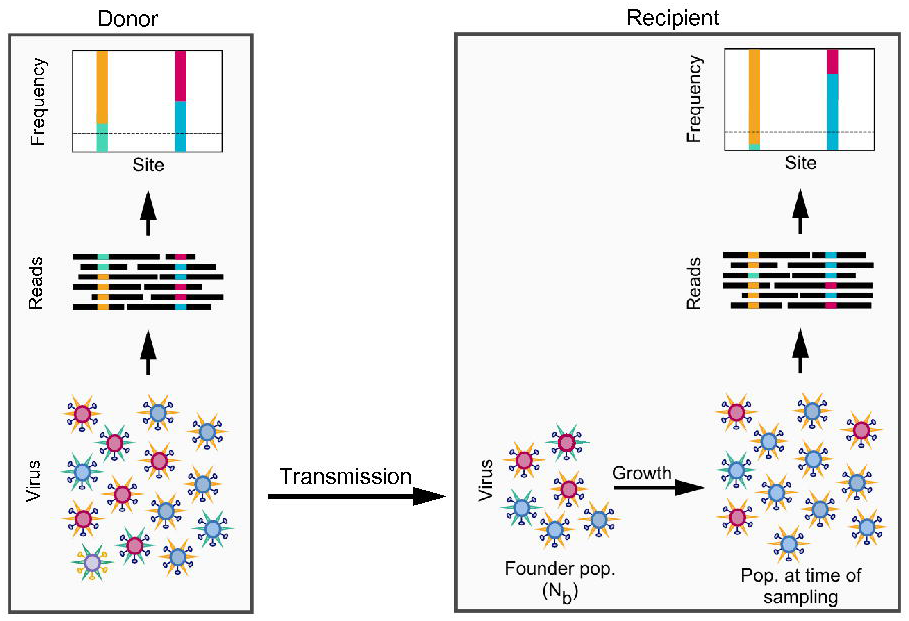
Schematic showing virus transmission from donor to recipient host. The number of virions that initiate infection in the recipient host is defined as the transmission bottleneck size or founding population size *N_b_*. The viral sampling process is shown, with deep sequencing of the viral population resulting in reads that carry polymorphisms at certain nucleotide sites. The nucleotide read-outs at any site can be used to estimate variant frequencies. Dashed horizontal lines in the variant frequency plots denote the variant calling cutoff or threshold. The goal is to estimate *N_b_* given data on variant frequencies in the donor, and in the recipient, the total number of reads and the number of variant reads at each of the variant sites identified in the donor.

Given the extent of sequencing error in deep-sequencing data, there can be a high degree of noise in the short read data and, thereby, in the extent of polymorphism present at nucleotide sites. To limit spurious identification of variants arising from sequencing noise, it is common practice to use criteria, such as a variant calling threshold, to validate identified variants [31]. The variant calling threshold is the minimum frequency at which a variant can almost certainly be distinguished from background sequencing error. This threshold frequency may be chosen according to generally accepted error rates for a specific sequencing platform, error rates informed by a control run, or error rates based on the concordance of variant calls from replicate sequence runs. For the commonly used Illumina sequencing platforms, variant thresholds tend to fall in the range of 0.5−3% [24–26,28,32–35]. Conservative variant calling cutoffs are often used, as they ensure that sequencing artifacts are excluded. However, conservative frequency cutoffs may have effects on transmission bottleneck size analyses due to variants that are not called in the recipient host, despite being present. Such ‘false negatives’ in the recipient have the potential to skew inferred transmission bottleneck size towards inappropriately low values.

We present methods for inferring the transmission bottleneck size from deep sequencing data, paying special attention to the effects of ‘false negative’ variant calls. We first introduce the beta-binomial sampling method that we have developed for bottleneck size inference, which further incorporates the effects of stochastic pathogen dynamics in recipient hosts. For comparison, we then summarize two existing methods of bottleneck size inference in the literature: the presence/absence method and the binomial sampling method. Of note, all three of these methods assume that the genetic diversity of the pathogen is entirely neutral, such that selection does not impact variant frequency dynamics. These methods further assume independence between variant sites. We address the limitations of these assumptions in the Discussion.

## Bottleneck size inference allowing for stochastic pathogen dynamics in the recipient host

The beta-binomial sampling method for inferring the bottleneck size allows variant allele frequencies in the recipient host to change between the time of founding and the time of sampling (see Figure 1) as the result of stochastic pathogen replication dynamics early in infection. We consider two implementations of the beta-binomial sampling method: an approximate version that assumes infinite read depth and an exact version that incorporates sampling noise arising from a finite number of reads. The derivation of the beta-binomial sampling method can be found in the Methods.

In the approximate version, the likelihood of a transmission bottleneck size *N*_b_, given variant frequency data at site *i*, is given by:

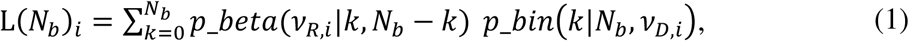

where *ν*_*R, i*_ is the variant frequency at site *i* in the recipient and *p_beta(V_R,i_|k, N_b_-k*) is given by the beta probability density function parameterized with shape parameters *k* and *N*_b_-k, and evaluated at *ν*_*R, i*_. The term *p_bin*(*k*|*N_b_,V_D,i_*) denotes the binomial distribution evaluated at *k* and parameterized with *N*_b_ number of trials and a success probability of *v*,*_D,i_*, where *v_D,i_* is the variant frequency at site *i* in the donor. If the donor variant at site *i* is not detected in the recipient, this may be because it is truly absent from the recipient or because it falls below the variant calling threshold. To allow for both of these possibilities, the likelihood that the transmission bottleneck size is *N*_b_, given that the variant at site *i* was not detected, is given by:

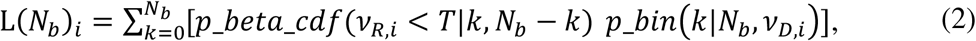

where *T* is the variant calling threshold and *p_beta_cdf*(*v_R,i_* < *T* | *k*, *N_b_ – k*) is given by the beta cumulative distribution function evaluated at the variant calling threshold.

In the exact version of the beta-binomial sampling method, we incorporate sampling error by modifying equations (1) and (2) to consider the number of variant reads and the number of total reads at variant site *i, R_var, i_* and *R_tot, i_*, respectively. The likelihood expression for the bottleneck size at site *i* becomes:

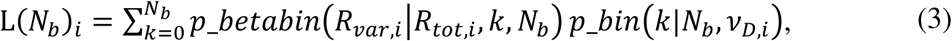

where *p_betabin*(*R_var,i_* \*R_tot,i_,k, N_b_*) is given by the beta-binomial probability density function evaluated at and *R_var,i_*.and parameterized with *R_tot,i_* number of trials and parameters *k* and *N*_b_. If the donor-identified variant at site *i* is not detected in the recipient, we again construct the likelihood that allows for this variant to either be absent from the recipient or below the variant calling threshold:

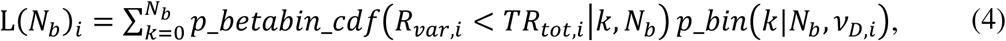

where, in this case, *p_betabin_cdf*(*R_var,i_* < *TR_tot,i_*|*k, N_b_*), is given by the beta-binomial cumulative distribution function evaluated at the number of reads that would qualify as falling at the variant calling threshold.

We expect that the maximum likelihood estimate (MLE) of *N*_b_ inferred with the approximate method will converge to the MLE of *N*_b_ inferred with the exact method when read coverage is high. The benefit of using the approximate version, when appropriate, is that the incorporation of sampling error is computationally intensive.

Once transmission bottleneck sizes have been estimated using either the approximate or exact beta-binomial sampling method, the probability that a variant is truly present/absent in the recipient and the probability that a variant is simply called present/absent in the recipient (under the assumption of infinite coverage) can be determined for any given donor variant frequency.

## Existing methods for inferring transmission bottleneck sizes

### Presence/absence method of bottleneck size inference

The simplest approach to estimating transmission bottleneck sizes from pathogen deep-sequencing data is to calculate variant frequencies in donor hosts and then use information on the presence/absence of these variants in recipient hosts to quantify bottleneck size. Studies that have adopted this approach include [9,36]. Given a variant *i* present at frequency *vD,i* in the donor, and a founding population size of *N*_b_, the probability that the variant was not transferred to the recipient is simply given by (1 − *v_D,i_*)^*N_b_*^ [9,36]. Correspondingly, the probability that at least one virion in the founding population carried the variant allele is given by 1 − (1 − *v_D,i_*)^*N_b_*^. From these expressions, the likelihood of the founding population size *N*_b_ in a donor-recipient pair is simply calculated by multiplying the probabilities of the observed outcomes across the variant sites:

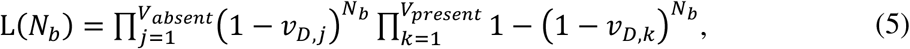

where *j* indexes the viral variants that are absent in the recipient, *k* indexes the viral variants that are present in the recipient, *V_absent_* is the total number of variants which are called absent in the recipient, and *V_present_* is the total number of variants that are called present in the recipient. The total number of variants identified in the donor is given by *V_absent_* + *V_present_*.

The presence/absence method considers only the detection of donor-identified variants in the recipient host and, therefore, is especially prone to the effects of false negative variants. Moreover, accounting for the variant calling threshold to ameliorate these effects is not possible with this method. Due to the inability of this method to account for false negatives, we expect that the transmission bottleneck estimates inferred with the presence/absence method will be considerably lower than the bottleneck size estimates inferred by the beta-binomial sampling method.

### Binomial sampling method of bottleneck size inference

The second approach, or class of approaches, from the literature for inferring transmission bottleneck sizes is based on a binomial sampling process. Studies that have adopted this general kind of approach include [28,30]. We describe a version of this approach that parallels the beta-binomial sampling method we described above. The binomial sampling approach makes use of donor-identified variant frequencies in the donor and both the number of variant reads and the number of total reads in the recipient, at each donor-identified variant site. The likelihood expression for the bottleneck size, given these data at site *i*, is given by:

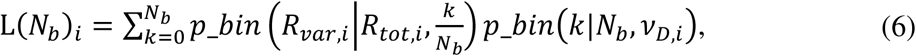

where 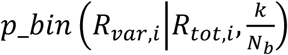 is given by the binomial probability density function evaluated at *R_var,i_*. The term *p_bin*(*k*|*N_b_,v_D,i_*) is again given by the binomial distribution. For variants called as absent in the recipient host, the likelihood of the transmission bottleneck size is given:

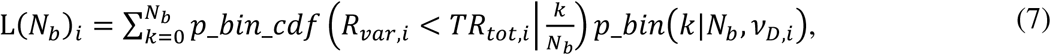

Where *p_bin_cdf* is the binomial cumulative distribution function. Derivation of the binomial sampling method can be found in the Methods section.

The sole difference between the beta-binomial sampling method and the binomial sampling method is that the binomial sampling method does not account for stochastic dynamics of the pathogen early on in the recipient. These stochastic dynamics enable the frequencies of variants in a recipient at the time of sampling to differ from those at the time of founding (Figure 1). Because the binomial sampling method does not incorporate this source of frequency variation, we expect there to be smaller frequency deviations between variants in donor-recipient pairs under the assumption of a single-generation binomial sampling model as compared to a model that allows for these stochastic dynamics, for a given bottleneck size. To explain a given pattern of donor-recipient frequency pairs, *N*_b_ estimates are thus expected to be significantly lower for the binomial sampling method than for the beta-binomial sampling method. Application of the binomial sampling method will therefore yield a conservative (lower-bound) estimate of *N*_b_, as previously remarked upon [30].

## Results

### Results on simulated data

To examine the ability of the three methods described above to accurately infer transmission bottleneck sizes, we used a simulated dataset of one donor-recipient pair (Methods). The dataset was generated under the assumption of stochastic pathogen dynamics in the recipient host between the time of infection and the time of sampling. While this assumption matches the assumption for the beta-binomial sampling method, we feel that it is also biologically the most realistic assumption. In this dataset, 109 out of the 500 donor-identified simulated variants were called absent in the recipient host (Figure 2A). The majority of these variants were present in the recipient host, but below our variant calling threshold of 3% and, therefore, were ‘false negatives’. The beta-binomial sampling method, as expected, recovers the true bottleneck size of 50 virions (Figure 2B). In contrast, both the presence/absence method (Figure 2C) and the binomial sampling method (Figure 2D) significantly underestimate the simulated bottleneck size. The underlying reasons for these methods’ inability to recover the true bottleneck size differ. For the presence/absence method, this underestimation can be attributed to ‘false negative’ variant calls. For the binomial sampling method, we were able to statistically account for the variant calling threshold effects; the underestimation of this method, therefore, is solely attributed to this method not accounting for stochastic pathogen dynamics in the recipient. The binomial sampling method instead assumes deterministic viral growth from the time of founding to the time of sampling (see Methods). Because more sampling stochasticity is present at smaller bottleneck sizes, the binomial sampling method underestimates the simulated bottleneck size in its attempt to reproduce observed variation in variant frequencies by inappropriately constricting *N*_b_.

**Figure 2.**
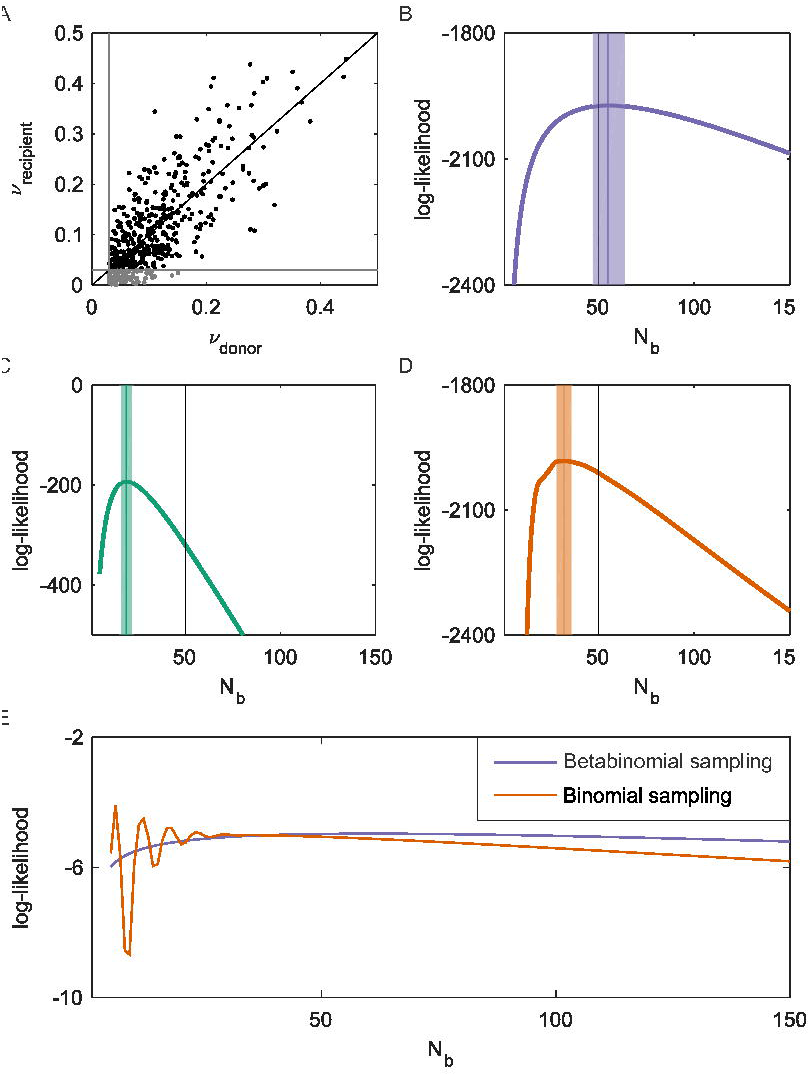
Estimated transmission bottleneck sizes for a simulated NGS dataset. (A) Scatterplot showing the frequencies of donor-identified variants against corresponding frequencies of these variants in the recipient. Points in black are variants that are called present in the recipient host. Points in grey are variants that are called absent in the recipient host. Black line shows where *ν*_donor_ = *ν*_recipient_. Gray lines show the variant calling threshold of 3%. (B) The beta-binomial sampling method’s log-likelihood curve over a range of *N_b_* values. Maximum likelihood estimate (MLE) = 55 virions (95% CI = 47-64 virions). Likelihood at MLE = -1972.7. (C) The presence/absence method’s log-likelihood curve over a range of *N_b_* values. MLE = 19 virions (95% CI = 16-22 virions). (D) The binomial sampling method’s log-likelihood curve over a range of *N_b_* values. MLE = 32 virions (95% CI = 28-36 virions). Likelihood at MLE = -1981.8. In (B)-(D), vertical black lines show the true transmission bottleneck size of *N_b_* = 50. Vertical colored lines show the MLE, and shaded areas show the 95% confidence interval, determined using the likelihood ratio test. Likelihood surfaces for a single variant present in the recipient at a frequency of 16.9% under the beta-binomial sampling model and the binomial sampling model.

Given that the binomial sampling model and the beta-binomial model fit to the same data, the relative performance of these models can be assessed using model selection approaches. The maximum likelihood obtained using the beta-binomial sampling method was significantly higher than the maximum likelihood obtained using the binomial sampling method (Figure 2B,D; legend), indicating that the beta-binomial sampling model is statistically preferred over the binomial sampling model. We can further take into consideration the smoothness of the likelihood curves in our choice of model, with multi-modal/rugged likelihood curves being undesirable outcomes. In Figure 2E, we plot the likelihood curves for one variant under the likelihood expression of the beta-binomial sampling method and under the expression of the binomial sampling method. The rugged likelihood surface of the binomial sampling model arises because of this method’s stringent assumption that variant frequencies remain fixed between the time of infection of the recipient and the time of sampling. In contrast, the beta-binomial sampling method allows for stochastic changes in variant frequencies during viral growth, relaxing the assumption that the viral population at the time of sampling needs to perfectly reflect the founding viral population. As a result, likelihood curves of the beta-binomial sampling model do not show large differences in likelihood values for small differences in *N*_b_, further indicating that the beta-binomial sampling model is preferable.

Given an estimate of the transmission bottleneck size, the probability that a variant is transferred to a recipient host can be calculated using the expression 1 − (1 − *v_D,i_*)^*N_b_*^, where *ν*,*_D,i_* is the frequency of variant *i* present in the donor host and *N*_b_ is the bottleneck size estimate. In Figure 3A, we plot this probability of variant transfer over a range of donor variant frequencies for the simulated dataset. In this figure, we further plot ‘observed’ probabilities of variant transfer, given a variant calling threshold of 3% on the simulated dataset. Finally, we plot in this figure the ‘observed’ probabilities of variant transfer as predicted under the beta-binomial sampling method, evaluated at the transmission bottleneck size estimated. We see, first, that the true probabilities of variant transfer greatly exceed those that are observed in the dataset given the variant calling threshold of 3%. However, the method’s calculated predictions of observed variant transfer probabilities fall within the 95% confidence intervals for the probabilities of variant transfer observed in the dataset.

**Figure 3.**
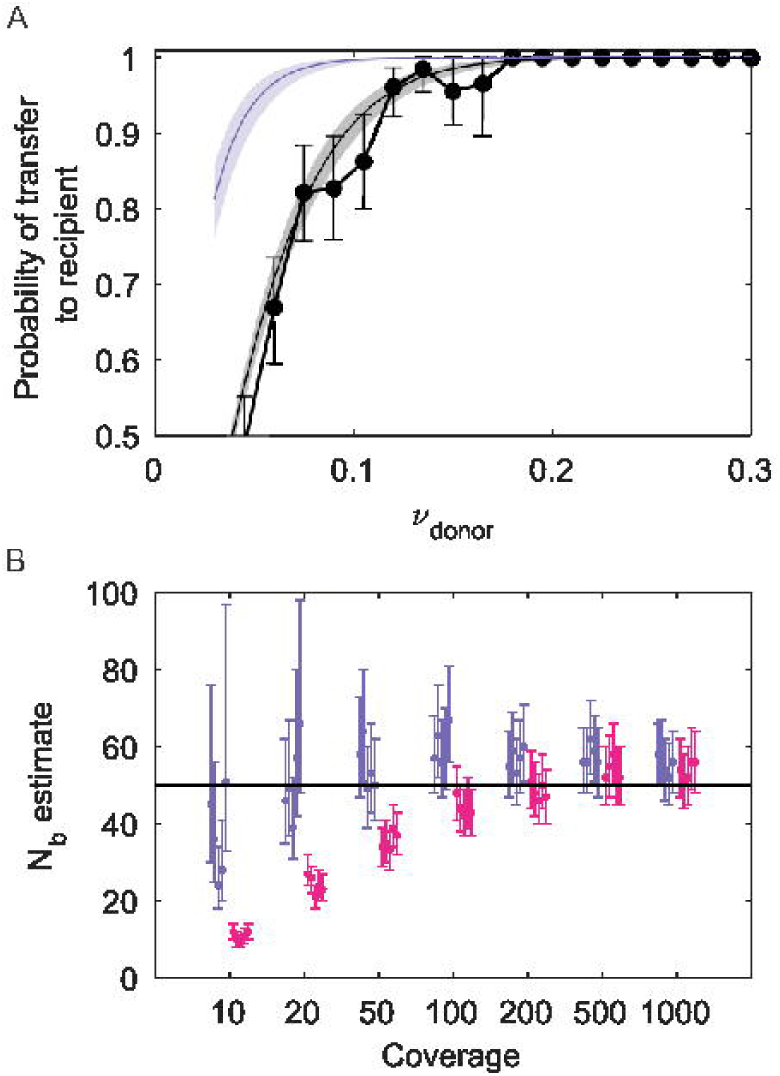
Additional results from application of the beta-binomial sampling method to thesimulated dataset. (A) The probability of a donor-identified variant being either transferred or observed as transferred (“called”) in a recipient host, as a function of donor variant frequencies. Observed probabilities of donor-identified variants being called in a recipient host are shown in black, calculated directly from the simulated dataset using 3% frequency bins. 95% confidence intervals assume the probability of variant transfer follows a binomial distribution with the number of trials being the number of donor-identified variants present in a frequency bin and the success probability given by the calculated probability of transferred variants observed in the frequency bin. Probabilities of donor-identified variants being truly present in a recipient host are shown in purple, given bottleneck size estimates from the beta-binomial sampling method. Probabilities of donor-identified variants being called present in a recipient host are shown in gray, given bottleneck size estimates from the beta-binomial sampling method. (B) *N_b_* estimates for simulated datasets that differ in coverage levels. At each coverage level, 5 datasets were generated, under the same parameters and assumptions as the dataset shown in Figure 2A. The exact beta-binomial sampling method and the approximate version of this method were both used to estimate *N_b_* for each dataset. *N*_b_ maximum likelihood estimates and 95% confidence intervals are shown, in purple for the exact beta-binomial sampling method and in pink for the approximate method.

As described in the Models section, the exact beta-binomial sampling method we developed accounts for sampling noise arising from finite read coverage. If we ignore sampling noise, we can estimate bottleneck sizes more rapidly using the approximate method, described by equations (1) and (2). In Figure 3B we show bottleneck size estimates over a range of different coverage levels for both the exact and approximate beta-binomial sampling methods. At high coverage levels (>200 reads), both implementations of the beta-binomial sampling method yield similar bottleneck size estimates and are able to recover the simulated bottleneck size of 50 virions. For lower levels of coverage, however, this approximation starts to fail and will lead to a considerable underestimation of *N*_b_, indicating that the approximate beta-binomial sampling method is inappropriate for low coverage levels. We also note that even at high coverage, a slight overestimation of the bottleneck size is apparent. The overestimation can be attributed to the rare false positive identification of variants in the recipient (instances of a variant that is absent in the recipient being called present) and, more generally, a slight inflation of variant frequencies with sequencing error. Overestimation no longer occurs when these methods are applied to datasets that are simulated in the absence of sequence error (results not shown).

### Transmission Bottleneck Size Estimation for Human Influenza A Virus

We first applied the beta-binomial sampling method for inferring transmission bottleneck sizes to the influenza A/H1N1p transmission pairs identified in a previously studied influenza NGS dataset described in detail in [28]. We point the reader to this previous publication for details on the dataset, including coverage levels, how transmission pairs were inferred, etc. Poon and Song et al. [28] estimated the mean effective population size for all H1N1p transmission pairs to be *N*_e_ = 192 virions (mean s.d. range 114-276). The approach considered the combined set of variants that were present at frequencies ≥1% and that were shared by 8 identified household donor-recipient pairs (a total of 26 variants). In contrast to their analysis, we estimated transmission bottleneck sizes for each of 9 transmission pairs separately, using a minimum variant frequency cutoff of 3% to call variants. We used a 3% cutoff based on concordance results from replicate sequencing runs, described in [28]. The less conservative 1% cutoff used by Poon and Song et al. [28] to estimate effective population size was chosen to allow for more sites to be included in their analysis. Our analysis, using a total of 289 variants, estimated MLE bottleneck sizes ranging from 49 to 276 virions across the H1N1p transmission pairs (Figure 4A). The bottleneck sizes inferred by the approximate beta-binomial sampling method did not differ significantly from those inferred by the exact method for any of the transmission pairs. This was expected, given high coverage levels across variant sites.

**Figure 4.**
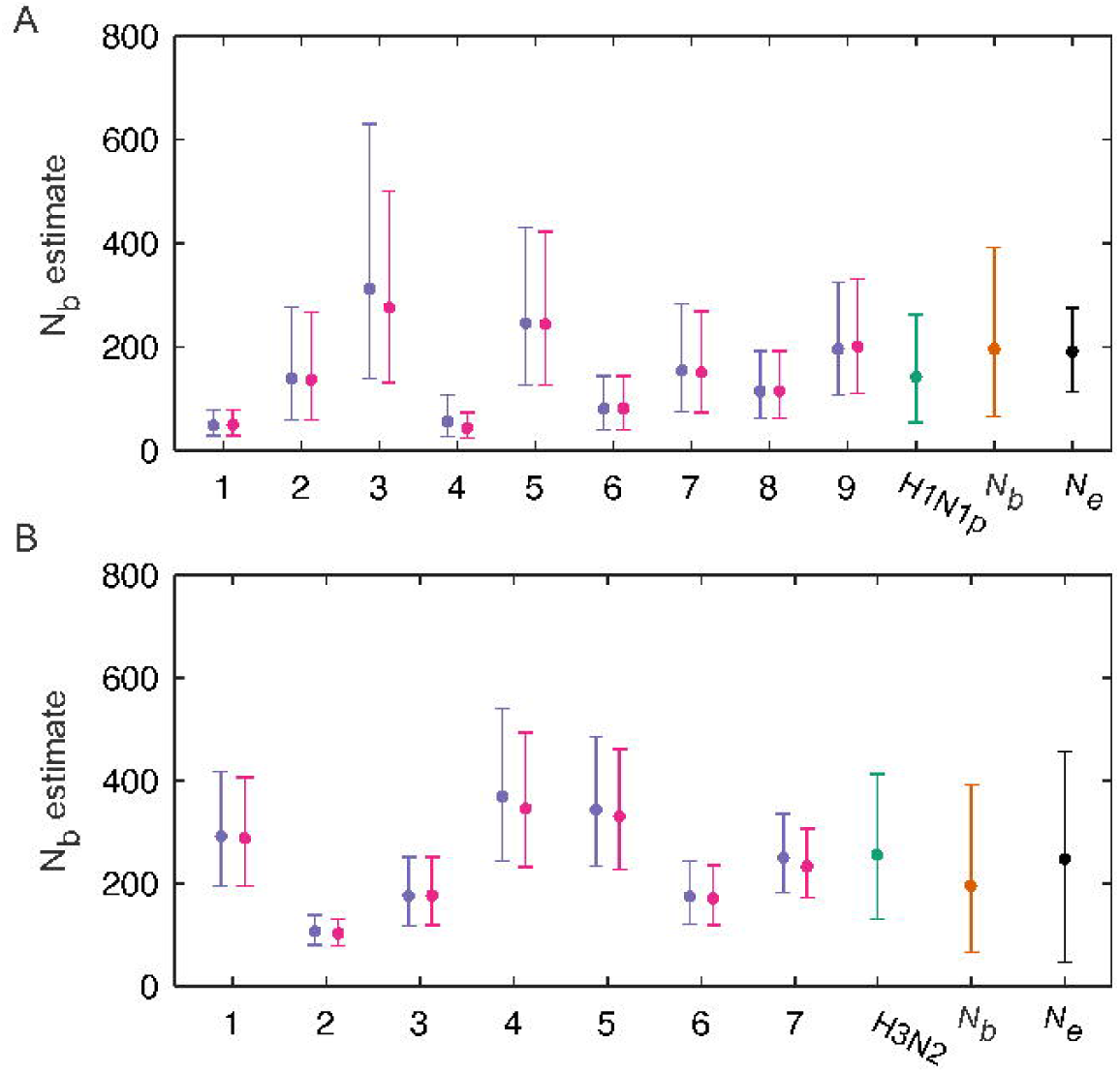
Transmission bottleneck sizes estimated for influenza A virus transmission pairsH1N1p (A) and H3N2 (B). *N_b_* estimates are shown for the exact beta-binomial sampling method (purple) and the approximate version of this method (pink). Bars show mean and 95% CI, calculated using the likelihood ratio test. Overall transmission bottleneck sizes estimated across H1N1p transmission pairs (‘*H1N1p*’, teal), across H3N2 transmission pairs (‘*H3N2*’, teal), and across both subtypes (‘*N_b_*’, orange), under the assumption of a negative binomial distribution, are also shown. Previous Poon et al. estimates are further shown (‘*N*e’) for H1N1p and H3N2 (black). Bars for the Poon et al. estimates show mean estimated effective population sizes and mean s.d. ranges.

To summarize our results for the bottleneck size estimates for the H1N1p transmission pairs, we estimated parameters of a negative binomial distribution using all of the variant frequencies across the transmission pairs (see Methods). This negative binomial distribution was chosen because our results shown in Figure 4A indicated that the variance in transmission bottleneck sizes is likely to exceed the mean. We further fit a Poisson distribution to these same data, and the negative binomial distribution was statistically preferred over the Poisson distribution using AIC, indicating that, while a single infection may be initiated by a Poisson-distributed number of virions, different infections are likely to be initiated by founding population sizes that vary in their mean. The MLE of the negative binomial distribution’s parameters was *r* = 5 and *p* = 0.966, resulting in a mean H1N1p transmission bottleneck size of *N*_b_ = 142, and a 95% range of 54-262 virions (Figure 4A). While our overall bottleneck size estimates were consistent with the estimates from Poon and Song et al. using a much more limited number of variants, our analysis further shows that the transmission bottleneck sizes varied considerably between transmission pairs.

We next used the beta-binomial sampling method to infer the transmission bottleneck sizes for each of the H3N2 transmission pairs of the influenza NGS dataset. Poon and Song et al. estimated the mean effective population size for H3N2 to be *N*_e_ = 248 (mean s.d. range 45-457), again using a combined set of variants that were present at frequencies ≥1% and that were shared by 6 identified household donor-recipient pairs (a total of 81 variants). Our analysis, considering each of the 7 identified H3N2 transmission pairs separately, inferred MLE bottleneck sizes ranging from 107 to 370 virions across the transmission pairs using a total of 621 variants (Figure 4B). Again, as expected, the *N*_b_ sizes inferred by the approximate beta-binomial sampling method did not differ significantly from those inferred using the exact beta-binomial sampling method. We again fit a negative binomial distribution to all of the variants across the transmission pairs and estimated MLE parameters of r = 9 and *p* = 0.966, resulting in a mean H3N2 transmission bottleneck size of *N*_b_ = 256, and a 95% range of 131-413 virions (Figure 4B). We again observed that the overall bottleneck size estimate for H3N2 was consistent with Poon et al.’s estimate, though the bottleneck size estimates varied considerably between transmission pairs.

#### Overall influenza A transmission bottleneck sizes

We next sought to determine whether influenza A/H1N1p and influenza A/H3N2 virus subtypes statistically differed from one another in bottleneck sizes. We found that the H1N1p and H3N2 distributions of transmission bottleneck size MLEs did not differ significantly from one another (*p* = 0.15 using the Kolmogorov-Smirnov test). Given this finding, we fit a negative binomial distribution to all of the variants across both subtype datasets, arriving at a MLE of *r* = 4 and *p* = 0.980 for the parameters of the negative binomial distribution. These parameters correspond to a mean bottleneck size of *N*_b_ = 196 and a 95% range of 66-392 virions (Figures 4A, 4B). We show the probability density function for this negative binomial distribution in Figure 5A. We further plot the expected probability of variant transfer for this bottleneck size estimate (Figure 5B), similar to what we show for the simulated dataset in Figure 3A. Finally, we plot the probability of observed variant transfer under this *N*_b_ estimate, under the assumptions of the beta-binomial sampling model. The agreement between the probability of observed variant transfer and the empirical data indicate that variant calling thresholds again make it appear that variant transfer from donor to recipient is much less likely than it is, given bottleneck size estimates based on variant frequencies.

**Figure 5.**
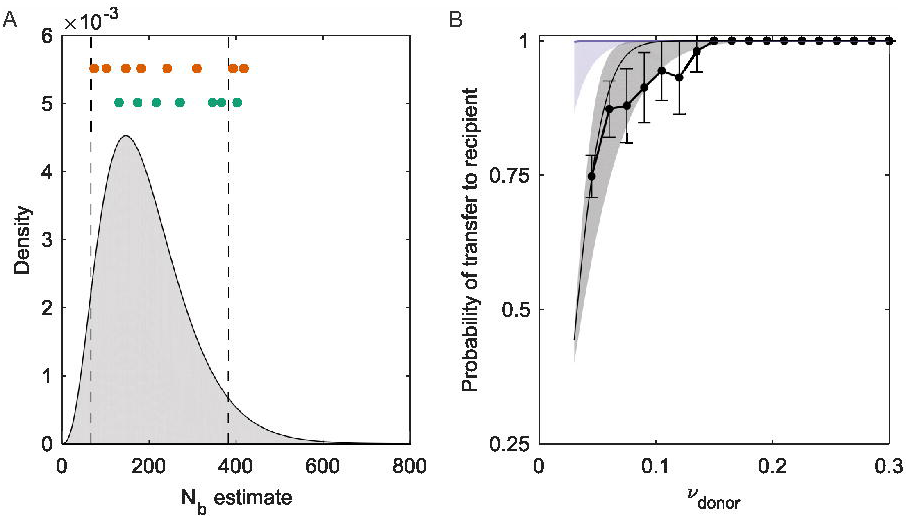
Overall influenza A bottleneck size estimates and probabilities of variant transfer under these estimates. (A) The negative binomial probability density function describing overall transmission bottleneck sizes across H1N1p and H3N2 viral subtypes, parameterized with the MLE values of *r* = 4 and *p* = 0.980. Vertical black lines show the 95% range of this distribution. The MLE bottleneck size estimates for the H3N2 (orange) and H1N1 (green) transmission pairs are shown above the pdf. (B) The probability of a donor-identified variant either being transferred or identified (“called”) in the recipient host, as a function of donor variant frequency. Probabilities of a donor variant being present in a recipient host are shown in purple, given bottleneck size estimates provided by the negative binomial distribution shown in (A). Probabilities of donor identified variants being called present in a recipient host, given these same bottleneck size estimates and the assumptions of the beta-binomial sampling models, are shown in gray. The empirical probabilities of donor-identified variants being called in a recipient, as calculated from the combined H1N1p and H3N2 datasets over 3% frequency bins, are shown in black.

## Discussion

Here, we have introduced a new method for estimating the transmission bottleneck size of pathogens from next generation sequencing data from donor-recipient pairs. We have further analyzed how well this beta-binomial sampling method performs in comparison to two existing methods in the literature: the presence/absence method and the binomial sampling method. Using a simulated dataset, we have demonstrated that both the presence/absence method and the binomial sampling method (for different reasons) systematically underestimate the transmission bottleneck size and that the latter can lead to undesirable rugged likelihood curves. In contrast, the beta-binomial sampling method, as expected, is able to recover the simulated bottleneck size (Figure 2B) and is able to accurately predict the probability that a donor variant would be identified in a recipient host under a given variant calling threshold (Figure 3A). Application of the beta-binomial sampling method to a previously published H1N1p and H3N2 NGS dataset showed a high degree of heterogeneity between bottleneck size estimates across transmission pairs (Figure 4). A negative binomial distribution was fit to all of the variants, yielding an overall mean *N*_b_ of 196 virions and a 95% range of 66 – 382 virions (Figures 4A, 4B, 5A).

The bottleneck sizes that we estimated for the H1N1p and H3N2 transmission pairs are close to the previous estimates of the effective population size *N*e arrived at by Poon and Song et al. for this dataset [28], although we were able to further estimate transmission bottleneck sizes by transmission pair and our method was able to make use of a much larger number of identified variants. Our bottleneck size estimates are consistent with the more qualitative observations of loose transmission bottlenecks for influenza A virus transmission in horses [20,22,23], pigs [20,21], and dogs [19]. Our *N*_b_ estimates, however, are considerably larger than the previous bottleneck sizes estimated for this virus by Varble et al. [27], Frise et al. [29], and McCaw et al. [17]. Varble et al.’s experimental study showed that the route of transmission affected the bottleneck size, with contact transmission giving rise to larger bottlenecks. They found that, of the 71-100 distinct viral tags, only 7-24 of those tagged viruses were detected in the recipients following infection via direct contact [27]. The number of distinct viral tags, however, might reflect the lower limit of the bottleneck size because it is possible that more than one virion passing through the bottleneck would have the same tag. Frise et al. reported a mean bottleneck size of 28.2 infectious genomes for contact transmission of an efficiently transmitted H1N1 strain in ferrets, though they were unable to identify an upper limit to the bottleneck size confidence interval [29]. Both of these previous estimates are much larger than the earlier estimate of 3.8 virions estimated by McCaw et al. for contact transmission of H1N1 in ferrets [17]. While other studies exist that have estimated the transmission bottleneck size in the context of viral adaptation to a new host species [24–26], comparisons with these studies are inappropriate because these bottlenecks are subject to strong selective forces, which considerably narrow the transmission bottleneck size [37].

The *N*_b_ estimates for influenza virus transmission in the dataset described here, both by our study and Poon and Song et al.’s original analysis, are considerably higher than previous quantitative estimates of IAV’s bottleneck size for contact transmission [17,27,29]. Notably, these previous estimates of *N*_b_ were arrived at using data from experimental ferret infections. With a recent analysis showing that secondary attack rates in ferret studies are considerably higher than human secondary attack rates, controlling for infecting subtype [38], one possibility for these discrepancy is that ferrets and other small mammals may require fewer influenza A virions to successfully initiate infection.

McCaw et al.’s bottleneck size estimate, in particular, was significantly lower than our *N*_b_ estimates for contact transmission of influenza virus [17]. One possible explanation for the low *N*_b_ estimate is that the ‘competitive mixture’ method they used to calculate bottleneck size considers only two viral populations, analogous to the estimates derived from a single variant in the methods we considered. The competitive mixture method is, thus, highly susceptible to fluctuations between donor and recipient variant frequencies arising from stochastic viral dynamics in the recipient. Thus, for the same reason that the binomial sampling method we described here underestimates bottleneck sizes, we would expect this competitive mixture method to considerably underestimate bottleneck sizes. Yet, this method is free of one of the necessary assumptions made for each the three methods that we considered, namely that the variants considered are independent. The independence assumption is clearly violated in this data set given extensive genetic linkage within influenza gene segments [39]. We can, however, somewhat control for the effects of linkage by selecting only one variant per gene segment. This data-thinning approach still assumes independence across gene segments that, while not ideal, may be supported by recent experimental evidence showing high levels of reassortment *in vitro* [40]. If intrahost reassortment occurs at similar rates *in vivo*, then sampling only one variant per gene segment should remove much of the bias due to linkage.

The methods we considered make other assumptions that may have also impacted transmission bottleneck size estimates. These assumptions include that: (1) donor-identified variants did not originate *de novo* in any recipient hosts, (2) variants were biallelic, and (3) variants were selectively neutral. Significant levels of *de novo* evolution of variants in recipient hosts would artificially increase estimated bottleneck sizes. Therefore, these methods may not be appropriate for pathogens causing chronic infections, such as HIV, where sampling of the recipient host can occur years after infection initiation. However, we do not expect substantial *de novo* evolution of variants to occur over the course of an acute influenza infection based on recent findings [37] and the observation that the vast majority of recipient-identified variants were also present in the donor. Therefore, we do not expect this assumption to have significantly influenced our bottleneck size estimates for influenza virus.

We also do not expect that the second assumption—that loci are biallelic—to have biased our bottleneck size estimates. This is because no sites used in our bottleneck size calculations contained more than one variant allele above our variant calling threshold of 3%. This assumption, however, could be removed in future uses of the beta-binomial sampling method by appropriately modifying the likelihood expressions to account for more than one variant per site.

The third assumption of selective neutrality is the one that could greatly affect the accuracy of our bottleneck size estimates if not met. Selection, either for or against a variant, would lead to larger differences in variant frequency between a donor and a recipient host than would be expected for neutral variants. Larger differences in variant frequencies would bias the estimated transmission bottleneck sizes towards smaller values. Thus, our bottleneck size estimates, which assume neutrality, are necessarily conservative estimates.

In this study, we have developed a new statistical approach that can be used to accurately infer transmission bottleneck sizes for acute viral infections, such as influenza, RSV, and norovirus, using NGS data from identified donor-recipient pairs. This beta-binomial sampling method accounts for the possibility of ‘false negatives’ variants that are not called as present due to necessary variant calling thresholds. The method further accounts for changes in variant frequencies between the time of recipient infection and the time of pathogen sampling from the recipient that arise due to stochastic replication dynamics early in infection. Given the importance of the transmission bottleneck size in regulating the rate of pathogen evolution at the level of the host population, estimation of the transmission bottleneck size is a necessary component in the analysis of pathogens important to public health. Though methods such as viral tagging to estimate the bottleneck size for experimental infections exist, these techniques are not applicable for natural infections. Hence this work provides a strong foundation for future estimation of bottleneck sizes from viral sequence data that, importantly, can be applied to clinical samples.

## Materials and Methods

### Development of the beta-binomial sampling method

Here, we derive the beta-binomial sampling method for inferring transmission bottleneck sizes from pathogen NGS data. The final likelihood expressions for this method are provided in equations (3) and (4). As described above, the method allows variant frequencies in the recipient host to change between infection and sampling (Figure 1) due to stochastic pathogen dynamics occurring during the process of replication. More concretely, early in the infection when there are only a small number of replicating virions, stochasticity in viral growth is expected to have a large effect. For a stochastic birth-death process with a constant birth rate λ and a constant death rate *μ*, the probability mass function for the viral population size originating from a single virion that successfully establishes is given by [41]:

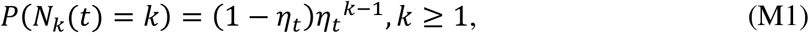

where *t* is the time of sampling and 
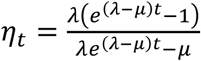
.For the bursty replication that characterizes many viruses, (M1) is still approximately true at long times with an adjusted value of η*_t_*.

The population sizes stemming from each of the *N_b_* founding virions, contingent on their successful establishment, are thus geometrically-distributed random variables. As these population sizes are likely to be very large at the time of sampling, we can approximate them as being exponentially-distributed random variables. Under this approximation, the distribution of the fractions of the population that descend from each of the founding virions is distributed as Dirichlet(1,1,…1), with *N*_b_ 1's, one for each ancestor. A subset *k* of these founder virions carry the variant allele; the remaining subset of these founder virions (*N*_b_-*k*) carry the reference allele. Collapsing the Dirichlet distribution yields that the fraction of the population carrying the variant allele is distributed as Beta(*k*, *N*_b_-*k*). Remarkably, this fraction does not depend on the within-host viral birth rate λ, the death rate *μ*, the time of sampling *t*, or the burstiness of replication. To get the overall likelihood of population bottleneck size *N*_b_, we simply have to consider all possible scenarios of how many virions out of the total *N*_b_ virions transferred carried the variant allele. Under the assumption that the founding pathogen population is randomly sampled from the pathogen population of the donor host, the probability that the founding population of *N*_b_ virions carries *k* variant alleles is given by the binomial distribution *p_bin*(*k*|*N_b_*, *v_D,i_*) ≡ 
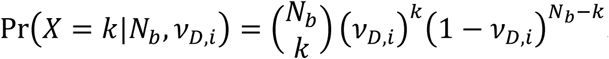
, where the number of trials is given by *N*_b_ and the success probability is given by *v_D,i_*, the frequency of variant *i* in the donor. Thus, the overall likelihood of population bottleneck size *N*_b_ for variant *i* is given by:

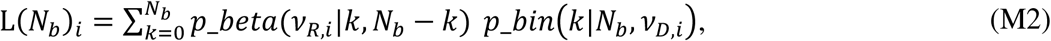

where *ν*_*R,i*_ is the frequency of variant *i* in the recipient and the term *p_bea*(*v_R,i_*|*k,N_b_-k*) is given by the beta probability density function, evaluated at *ν*_*R,i*_. This expression is provided in the main text as equation (1).

Accommodating sampling noise arising from a finite number of reads is simple, leading to minor modifications to the above equation (M2):

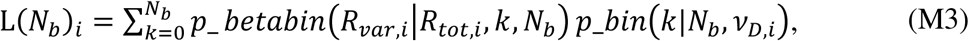

where *R_var,i_* is the number of reads of the variant allele in the recipient sample at site *i*, and *R_tot_* is the total number of reads at that site. The term *p_batabin*(*R_var,i_*|*R_tot,i_*, *k*, *N_b_*) is given by the beta-binomial distribution evaluated at *R_var,i_* and parameterized with *R_tot,i_* as number of trials and parameters *k* and *N*_b_. Expression (M3), reproduced in the main text as equation (3), thus incorporates both noise from the sampling process itself and from the process of stochastic pathogen growth. The overall likelihood of bottleneck size *N*_b_ for a transmission pair is simply the product of the site-specific likelihoods.

As previously mentioned, we expect that variant calling thresholds will impact the likelihood calculations used in the bottleneck size estimation. These thresholds will force some variant alleles in the recipient viral population to be called absent when they are actually present at frequencies below the value of the chosen threshold. Since true absence of a variant allele is more likely at smaller bottleneck sizes, conservative variant calling thresholds will bias *N*_b_ estimates to lower values. Simply excluding variants that are called absent from the analysis, however, will also bias bottleneck size estimates, this time towards higher values. To get around this, we do not recommend simply lowering the variant calling threshold because NGS sequencing error can give rise also to false positives, thereby inappropriately inflating bottleneck size estimates. Instead, we recommend accommodating below-threshold variants in the following way. For a donor-identified variant *i* that is called absent in the recipient (whether truly absent or just called absent), the likelihood of the transmission bottleneck size is given by the following expression:

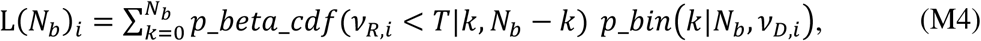

where *T* is the variant calling threshold (e.g., of 3%) and *p_beta_cdf*(*ν_R,i_* < *T*|*k, N_b_*−*k*) is given by the beta cumulative distribution function evaluated at the variant calling threshold. This expression is reproduced in the main text as equation (2). We can again incorporate the effects of sampling noise by considering the number of reads at the variant site with the expression:

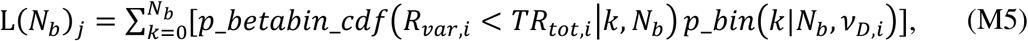

where, in this case, *p_betabin_cdf*(*R_var,j_* < *TR_tot,j_*|*k,N_b_*) is given by the beta-binomial cumulative distribution function evaluated at the number of reads that would qualify as falling at the variant calling threshold. This expression reproduces equation (4) of the main text.

Once the transmission bottleneck sizes have been estimated using the beta-binomial sampling method, the probability of true variant presence/absence in the recipient host can be determined for any given donor variant frequency. Similarly, the probability that a variant is called present/absent can be determined for any given donor frequency *ν*_*D,i*_, given a sufficiently high read count in the recipient host. Given a high read count, the probability that a variant is called present in the recipient is given by: 
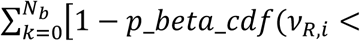
.

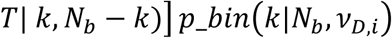

### The binomial sampling method

In contrast to the beta-binomial sampling method, the binomial sampling method implicitly assumes that the infecting virus population is subject to deterministic dynamics between the time of infection and the time at which the recipient virus is sampled and, thus, that the sampled pathogen population in the recipient perfectly reflects the founding pathogen population under the common assumption of selective neutrality. The founding pathogen population is, as in the beta-binomial sampling method, assumed to be randomly sampled from the pathogen population of the donor host. The site-specific likelihood of the transmission bottleneck size *N*_b_ is therefore given by:

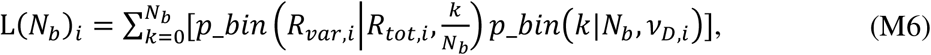

where 
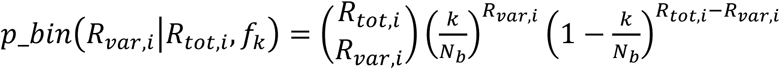
. This expression reproduces equation (6) in the main text. The overall likelihood of transmission bottleneck size *N*_b_ is calculated by multiplying across all site-specific likelihoods.

The above expression incorporates sampling noise, which is important when only a small number of reads are available. With an increasing number of reads, sampling noise necessarily goes down, making 
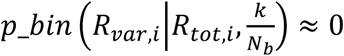
 in cases where 
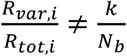
. This will result in dramatic differences in likelihood values between small values of *N*_b_, and more generally, multi-modal likelihood curves that are very sensitive to specific variant frequencies in the recipient host.

One basic issue with this approach is therefore the assumption of where differences in variant frequencies across donor-recipient pairs stem from. Under this model, any observed differences are due to the presence of a transmission bottleneck because it assumes that the sampled pathogen population in the recipient perfectly reflects the founding pathogen population. This assumption is met under a scenario of deterministic, and neutral, viral population dynamics between the time of the transmission event and the time of pathogen sampling from the recipient host. For example, if we assume deterministic exponential growth from the time of the transmission event to the time of sampling, the dynamics of the viral population that carries the variant allele is given by *N_ν_*(*t*) = *N*_ν_(0)*e^rt^*, and, similarly, the dynamics of the viral population that carries the reference allele is given by *N_r_*(*t*) = *N_r_*(0)*e^rt^*. At the time of the transmission event (*t* = 0), the fraction of the viral population that carries the variant allele is given by *k*/*N_b_*. At time, *t*, the fraction of the viral population that carries the variant allele is given by *N_v_*(*t*)/(*N_v_*(*t*) + *N_r_*(*t*)), which simplifies *k*/*N*_b_.

The bottleneck size estimates inferred with the binomial sampling method are again subject to the effects of ‘false negative’ variant calls. We can modify the binomial sampling method to incorporate the variant call threshold in a way similar to how the threshold frequency was incorporated into the beta-binomial sampling method. For a donor-identified variant *i* that is called absent in the recipient (whether truly absent or just called absent), the likelihood of the transmission bottleneck size is:

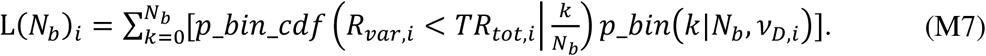

This expression reproduces equation (7) in the main text. The probability that the number of variant reads falls below the level required for the variant to be called present is given by the binomial cumulative distribution function:

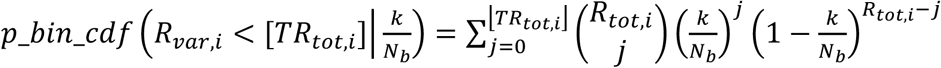

where ⌊*TR_tot,i_*⌋ is the largest integer smaller than *TR_tot,i_*.

As with the beta-binomial sampling method, once transmission bottleneck sizes have been estimated using the binomial sampling method, the probability of true variant presence/absence in the recipient host can be determined for any given donor variant frequency. Similarly, the probability that a variant is called present/absent can be determined for any given donor frequency *ν*_*D,i*_, provided information on the total read count in the recipient. Specifically, in the case of a high number of reads, the probability that a variant is called present (whether it is absent or present in the recipient host) is given by 
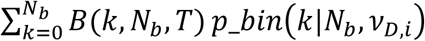
, where *B*(*k,N_b_,T*) is a Boolean function that evaluates to 1 if 
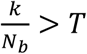
and 0 otherwise.

### Simulated deep-sequencing data

To illustrate the use of the methods used to estimate *N*_b_, we generated a mock deep-sequencing dataset via simulation. For this dataset, we assumed a single donor-recipient pair, with 500 independent donor-identified variants. Independently for both the donor and the recipient, we drew the total number of reads at each of the 500 sites from a normal distribution with a mean of 500 reads and a standard deviation of 100 reads. Draws from the normal distribution were rounded to the nearest integer and those that fell at 0 or below were discarded. For the donor, we then first determined “true” variant frequencies at each of these sites by drawing from an exponential distribution with mean frequency of 0.08. Variants with observed frequencies below the variant calling threshold of 0.03 or above 0.50 were discarded. To determine the number of variant reads at a given site in the donor, we drew from a binomial distribution with the number of trials being the total read count at that site in the donor and the probability of success being given by that site’s “true” variant frequency in the donor. We then incorporated sequencing error by again using draws from binomial distributions. Specifically, we determined the number of “true” reference reads in the donor that were misclassified as variant reads and the number of “true” variant reads in the donor that were correctly classified as variant reads, based on an assumed sequencing error rate of 1%. The total number of observed variant reads at a given site in a donor was then calculated as the sum of the misclassified reference reads and the correctly classified variant reads. Observed variant frequencies in the donor were then calculated by dividing the number of observed variant reads by the total number of observed reads at each site. In this manner, we simulated 500 variants, with observed frequencies in the range of 3-50%. The lower bound value of 3% was our assumed variant calling threshold; the upper bound value of 50% coincided with a variant allele always being the minority allele.

For the recipient, we simulated the total number of variant reads at each site by first simply determining at each site the number of virions in the founding population that carried the variant allele, under the assumption of a transmission bottleneck size of *N*_b_ = 50. This was done by, at each site, drawing from a binomial distribution with the number of trials being *N*_b_ and the probability of success being the “true” variant frequency at that site in the donor. For the simulated data set, we first determined the “true” fraction of the viral population carrying the variant allele at the time of sampling by drawing from a beta distribution with the shape parameter being the number of variant alleles in the founder population and the scale parameter being the difference between the founding population size of *N*_b_ and the number of variant alleles in the founder population. The “true” number of variant reads was then determined by drawing from a binomial distribution with the number of trials being the total number of reads at that site and the probability of success being the fraction of the population at the time of sampling that carried the variant allele. We then obtained the total number of variant reads at a given site in a recipient by introducing sequencing error to the “true” number of variant reads and the “true” number of reference reads.

### Application to Influenza A deep sequencing data

We applied the three methods for bottleneck size inference described in in the *Models* section to the influenza A deep-sequencing data examined in [28]. In this study, Poon and colleagues identified donor-recipient transmission pairs based on household information and the genetic similarities between the viral populations in infected hosts. We base our analyses on these already-identified transmission pairs. In some cases, there were several members of the household who became infected. In this subset of cases, rather than considering all feasible pairwise combinations of who-infected-whom, we assumed that the index case transmitted to multiple household members. With this assumption, the 9 identified transmission pairs for influenza A subtype H1N1p were 681_V1(0) → 681_V3(2), 684_V1(0) → 684_V2(3), 712_V1(0) → 712_V1(4), 742_V1(0) → 742_V3(3), 751_V1(0) → 751_V3(1), 751_V1(0) → 751_V2(3), 751_V1(0) → 751_V2(4), 779_V1(0) → 779_V2(1), 779_V1(0) → 779_V1(2), where *X*_V*Y*(*Z*) refers to household number *X*, visit number *Y*, subject *Z*, and the arrow demarcates transmission from donor to recipient. The 7 identified transmission pairs for influenza A subtype H3N2 were 689_V1(0) → 689_V2(2), 720_V1(0) → 720_V2(1), 734_V1(0) → 734_V3(2), 739_V1(0) → 739_V2(2), 739_V1(0) → 739_V2(3), 747_V1(0) → 747_V2(2), and 763_V1(0) → 763_V2(3). The deep-sequencing data are publically available from [28] and from https://www.synapse.org/#!Synapse:syn8033988. We called variants and determined variant frequencies from these data using VarScan [42,43], using a variant calling threshold of 3%, mean quality score of 20, and a p-value of 0.05. We provide variants and their frequencies as used in this study as a supplementary data file.

#### Calculation of overall transmission bottleneck sizes across transmission pairs

To calculate transmission bottleneck sizes over multiple transmission pairs, we did not simply take the sum of log-likelihoods across transmission pairs. Taking simply the sum would inappropriately give greater weight to transmission pairs with a larger number of donor-identified variants. To weight each of the transmission pairs equally, we scaled the log-likelihood of each transmission pair based on the number of variants identified in that transmission pair, such that the overall log-likelihood was given by 
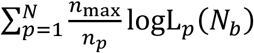

_h8_ abcd logL_h_, a9 where *N* is the number of transmission pairs, *n_p_* is the number of donor-identified variants in transmission pair *p*, *n_max_* = max(*n_p_*), and logL_*p*_ are the log-likelihoods across *N*_b_ values in transmission pair *p*.

## Acknowledgements

This work was funded by MIDAS CIDID U54-GM111274, supporting KK and ASL. ASL was further supported by the Duke Medical Scientists Training Program grant T32 GM007171. EG was supported by U01 AI111598.

## Bibliography

1. GutiérrezS, MichalakisY, BlancS.Virus population bottlenecks during within-host progression and host-to-host transmission. Curr Opin Virol. 2012;2: 546–555. doi:10.1016/j.coviro.2012.08.001

2. GeogheganJL, SeniorAM, HolmesEC. Pathogen population bottlenecks and adaptive landscapes: overcoming the barriers to disease emergence. Proc R Soc B Biol Sci. 2016;283: 20160727. doi:10.1098/rspb.2016.0727.

3. SkumsP, BunimovichL, KhudyakovY., Antigenic cooperation among intrahost HCV variants organized into a complex network of cross-immunoreactivity. Proc Natl Acad Sci U S A. 2015;112: 6653–8. doi:10.1073/pnas.1422942112

4. XueKS, HooperKA, OllodartAR, DingensAS, BloomJD. Cooperation between distinct viral variants promotes growth of h3n2 influenza in cell culture. Elife. 2016;5: 1–15.doi:10.7554/eLife.13974

5. BrookeCB, InceWL, WrammertJ, AhmedR, WilsonPC, BenninkJR, et al.Most influenza a virions fail to express at least one essential viral protein. J Virol. 2013;87: 3155–3162. doi:10.1128/JVI.02284-12

6. WorbyCJ, LipsitchM, HanageWP., Within-Host Bacterial Diversity Hinders Accurate Reconstruction of Transmission Networks from Genomic Distance Data. PLoS Comput Biol. 2014;10. doi:10.1371/journal.pcbi.1003549

7. De MaioN, WuC-H, WilsonDJ. SCOTTI: Efficient Reconstruction of Transmission within Outbreaks with the Structured Coalescent. 2016; Available: http://arxiv.org/abs/1603.01994

8. HallJS, FrenchR, HeinGL, MorrisTJ, StengerDC. Three distinct mechanisms facilitate genetic isolation of sympatric wheat streak mosaic virus lineages. Virology. 2001;282: 230–6. doi:10.1006/viro.2001.0841

9. SacristánS, MalpicaJM, FraileA, García-ArenalF., Estimation of Population Bottlenecks during Systemic Movement of Tobacco Mosaic Virus in Tobacco Plants. J Virol. 2003;77: 9906–9911. doi:10.1128/JVI.77.18.9906-9911.2003

10. MouryB, FabreF, SenoussiR.Estimation of the number of virus particles transmitted by an insect vector. Proc Natl Acad Sci U S A. 2007;104: 17891–6. doi:10.1073/pnas.0702739104

11. BetancourtM, FereresA, FraileA, García-ArenalF.Estimation of the effective number of founders that initiate an infection after aphid transmission of a multipartite plant virus. J Virol. 2008;82: 12416–12421. doi:10.1128/JVI.01542-08

12. ZwartMP, DaròsJA, ElenaSF. One is enough: In vivo effective population size is dose-dependent for a plant RNA virus. PLoS Pathog. 2011;7. doi:10.1371/journal.ppat.1002122

13. FabreF, MouryB, JohansenEI, SimonV, JacquemondM, SenoussiR., Narrow Bottlenecks Affect Pea Seedborne Mosaic Virus Populations during Vertical Seed Transmission but not during Leaf Colonization. PLoS Pathog. 2014;10: e1003833. doi:10.1371/journal.ppat.1003833

14. SmithDR, AdamsAP, KenneyJL, WangE, WeaverSC., Venezuelan equine encephalitis virus in the mosquito vector Aedes taeniorhynchus: Infection initiated by a small number of susceptible epithelial cells and a population bottleneck. Virology. 2008;372: 176–186. doi:10.1016/j.virol.2007.10.011

15. ZwartMP, HemerikL, CoryJS, de VisserJAGM, BianchiFJJ a, Van OersMM, et al.An experimental test of the independent action hypothesis in virus-insect pathosystems. Proc Biol Sci. 2009;276: 2233–2242. doi:10.1098/rspb.2009.0064

16. van der WerfW, HemerikL, VlakJM, ZwartMP., Heterogeneous host susceptibility enhances prevalence of Mixed-Genotype Micro-Parasite infections. PLoS Comput Biol. 2011;7. doi:10.1371/journal.pcbi.1002097

17. McCawJM, ArinaminpathyN, HurtAC, McVernonJ, McLeanAR. A Mathematical Framework for Estimating Pathogen Transmission Fitness and Inoculum Size Using Data from a Competitive Mixtures Animal Model. PLoS Comput Biol. 2011;7. doi:10.1371/journal.pcbi.1002026

18. ForresterNL, GuerboisM, SeymourRL, SprattH, WeaverSC., Vector-Borne Transmission Imposes a Severe Bottleneck on an RNA Virus Population. PLoS Pathog. 2012;8: e1002897. doi:10.1371/journal.ppat.1002897

19. HoelzerK, MurciaPR, BaillieGJ, WoodJLN, MetzgerSM, OsterriederN, et al.Intrahost Evolutionary Dynamics of Canine Influenza Virus in Naïve and Partially Immune Dogs. J Virol. 2010;84: 5329–5335. doi:10.1128/JVI.02469-09

20. StackJC, MurciaPR, GrenfellBT, WoodJLN, HolmesEC., Inferring the inter-host transmission of influenza A virus using patterns of intra-host genetic variation. Proc R Soc B Biol Sci. 2012;280: 20122173–20122173. doi:10.1098/rspb.2012.2173

21. MurciaPR, HughesJ, BattistaP, LloydL, BaillieGJ, Ramirez-GonzalezRH, et al.Evolution of an Eurasian Avian-like Influenza Virus in Naïve and Vaccinated Pigs. PLoS Pathog. 2012;8: e1002730. Available: http://www-ncbi-nlm-nih-gov.proxy.lib.duke.edu/pmc/articles/PMC3364949/?tool=pmcentrez&report=abstract.

22. HughesJ, AllenRC, BaguelinM, HampsonK, BaillieGJ, EltonD, et al.Transmission of Equine Influenza Virus during an Outbreak Is Characterized by Frequent Mixed Infections and Loose Transmission Bottlenecks. PLoS Pathog. 2012;8: e1003081. doi:10.1371/journal.ppat.1003081

23. MurciaPR, BaillieGJ, DalyJ, EltonD, JervisC, MumfordJA, et al.Intra and Interhost Evolutionary Dynamics of Equine Influenza Virus. J Virol. 2010;84: 6943–6954. doi:10.1128/JVI.00112-10

24. WilkerPR, DinisJM, StarrettG, ImaiM, HattaM, NelsonW, et al.Selection on hemagglutinin imposes a bottleneck during mammalian transmission of reassortant H5N1 influenza viruses. Nat Commun. 2013;4. doi:10.1038/ncomms3636.Selection

25. ZaraketH, BaranovichT, KaplanBS, CarterR, SongM-S, PaulsonJC, et al.Mammalian adaptation of influenza A(H7N9) virus is limited by a narrow genetic bottleneck. Nat Commun. Nature Publishing Group; 2015;6: 6553. doi:10.1038/ncomms7553

26. MonclaLH, ZhongG, NelsonCW, DinisJM, MutschlerJ, HughesAL, et al.Selective Bottlenecks Shape Evolutionary Pathways Taken during Mammalian Adaptation of a 1918-like Avian Influenza Virus. Cell Host Microbe. Elsevier Inc.; 2016;19: 169–180. doi:10.1016/j.chom.2016.01.011

27. VarbleA, AlbrechtRA a, BackesS, CrumillerM, BouvierNMM, SachsD, et al.Influenza A virus transmission bottlenecks are defined by infection route and recipient host. Cell Host Microbe. 2014;16: 691–700. doi:10.1016/j.chom.2014.09.020

28. PoonLLM, SongT, RosenfeldR, LinX, RogersMB, ZhouB, et al.Quantifying influenza virus diversity and transmission in humans. Nat Genet. Nature Publishing Group; 2016;1: 1–6. doi:10.1038/ng.3479

29. FriseR, BradleyK, van DoremalenN, GalianoM, ElderfieldRA, StilwellP, et al. Contact transmission of influenza virus between ferrets imposes a looser bottleneck than respiratory droplet transmission allowing propagation of antiviral resistance. Sci Rep. Nature Publishing Group; 2016;6: 29793. doi:10.1038/srep29793

30. EmmettKJ, LeeA, KhiabanianH, RabadanR., High-resolution Genomic Surveillance of 2014 Ebolavirus Using Shared Subclonal Variants. PLoS Curr. 2015;7: 1–17. doi:10.1371/currents.outbreaks.c7fd7946ba606c982668a96bcba43c90

31. OlsonND, LundSP, ColmanRE, FosterJT, SahlJW, SchuppJM, et al.Best practices for evaluating single nucleotide variant calling methods for microbial genomics. Front Genet. 2015;6: 235. doi:10.3389/fgene.2015.00235

32. GhedinE, HolmesEC, DePasseJ V., PinillaLT, FitchA, HamelinM-E, et al.Presence of Oseltamivir-Resistant Pandemic A/H1N1 Minor Variants Before Drug Therapy With Subsequent Selection and Transmission. J Infect Dis. 2012;206: 1504–1511. doi:10.1093/infdis/jis571

33. Van den HoeckeS, VerhelstJ, VuylstekeM, SaelensX., Analysis of the genetic diversity of influenza A viruses using next-generation DNA sequencing. BMC Genomics. 2015;16. doi:10.1186/s12864-015-1284-z

34. LakdawalaSS, JayaramanA, HalpinRA, LamirandeEW, ShihAR, StockwellTB, et al.The soft palate is an important site of adaptation for transmissible influenza viruses. Nature Nature Publishing Group, a division of Macmillan Publishers Limited. All Rights Reserved.; 2015;526: 122–5. doi:10.1038/nature15379

35. DinisJM, FlorekNW, FatolaOO, MonclaLH, MutschlerJP, CharlierOK, et al.Deep sequencing reveals potential antigenic variants at low frequency in influenza A-infected humans. J Virol. 2016;2013: JVI.03248–15. doi:10.1128/JVI.03248-15

36. SacristánS, DíazM, FraileA, García-ArenalF., Contact transmission of Tobacco mosaic virus: a quantitative analysis of parameters relevant for virus evolution. J Virol. 2011;85: 4974–4981. doi:10.1128/JVI.00057-11

37. Sobel LeonardA, McClainMT, SmithGJD, WentworthDE, HalpinRA, LinX, et al.Deep Sequencing of Influenza A Virus from a Human Challenge Study Reveals a Large Founder Population Size and Rapid Intrahost Viral Evolution. J Virol. 2016;

38. BuhnerkempeMG, GosticK, ParkM, AhsanP, BelserJA, Lloyd-SmithJO. Mapping influenza transmission in the ferret model to transmission in humans. Elife. 2015;4. doi:10.7554/eLife.07969

39. BoniMF, ZhouY, TaubenbergerJK, HolmesEC., Homologous Recombination Is Very Rare or Absent in Human Influenza A Virus. J Virol. 2008;82: 4807–4811. doi:10.1128/JVI.02683-07

40. InceWL, Gueye-MbayeA, BenninkJR, YewdellJW., Reassortment Complements Spontaneous Mutation in Influenza A Virus NP and M1 Genes To Accelerate Adaptation to a New Host. J Virol. 2013;87: 4330–4338. doi:10.1128/JVI.02749-12

41. KendallDG. On the Generalized “Birth-and-Death” Process. Ann Math Stat. 1948;19: 1–15.

42. KoboldtDC, ChenK, WylieT, LarsonDE, McLellanMD, MardisER, et al.VarScan: variant detection in massively parallel sequencing of individual and pooled samples. Bioinformatics. 2009;25: 2283–2285. doi:10.1093/bioinformatics/btp373

43. KoboldtDC, ZhangQ, LarsonDE, ShenD, McLellanMD, LinL, et al.VarScan 2: Somatic mutation and copy number alteration discovery in cancer by exome sequencing. Genome Res. 2012;22: 568–576. doi:10.1101/gr.129684.111

